# Single and Multi-Site Cortical Stimulation Related to Human Sensorimotor Function

**DOI:** 10.1101/2024.01.18.574786

**Authors:** Seokyun Ryun, Chun Kee Chung

## Abstract

Somatosensory feedback is crucial for precise control of our body and thereby affects various sensorimotor-related brain areas for movement control. Electrical stimulation on the primary somatosensory cortex (S1) elicits various artificial somatosensations. However, replicating the spatiotemporal dynamics of somatosensory feedback and fine control of elicited somatosensation are still challenging. Furthermore, how and where the somatosensory feedback interacts with neural activity for sensorimotor processing is unclear. Here, we replicate the spatiotemporal dynamics of somatosensory feedback and control the quality of elicited somatosensation using multi-site direct cortical stimulation (DCS). We also investigate how and where the neural feedback activity interacts with neural activity for motor processing by stimulating the downstream areas of the S1. We found that multi-site DCS on the S1 elicits different sensations simultaneously. Using the artificial feedback, blindfolded patients could efficiently perform a DCS-guided reach-and-grasp task successfully. Interestingly, we also found that multi-site DCS close to each other elicits different qualities of somatosensation in the same body part. Additionally, we found that DCS on the ventral premotor area (vPM) can affect hand grasping with eliciting artificial sensation of the hand. Throughout this study, we showed that semi-invasive, macro-level, and multi-site DCS can precisely elicit/modulate somatosensations in human. We suggest that activation of multiple cortical areas elicits simultaneous and independent somatosensations and that interplay among the stimulated sites can change the somatosensation quality. Finally, the results of vPM stimulation indicate that vPM has a critical role in function-specific sensorimotor interactions, such as hand grasping.

## Introduction

In everyday life, somatosensory feedback evoked by movement triggers neural orchestration of sensorimotor-related brain areas for precise movement control. Loss of somatosensory feedback causes critical impairments for precise movement and object manipulation (Augurelle et al., 2003; Sainburg et al., 1993), and the feedback information interacts with various sensorimotor-related brain areas for movement control (Omrani et al., 2016). Electrical brain stimulation is one of the most practical and widely adapted approaches to replicate such somatosensory feedback in humans (Flesher et al., 2016; Flesher et al., 2021). The findings that stimulating the appropriate area of the somatosensory homunculus evokes artificial somatosensation of specific body parts conceive the somatotopy concept (Penfield and Boldrey, 1937). A recent intracortical microstimulation (ICMS) study has successfully elicited somatosensation on a single body part with intra-digit precision (Flesher et al., 2016). Direct cortical stimulation (DCS) using high-density electrode grids showed spatial resolution as high as ICMS, although the elicited sensation’s naturalness was not tested yet (Kramer et al., 2021). Although the resolution of single-site stimulation has been sufficiently enhanced to elicit the artificial sensation in the exact area, our somatosensory feedback almost always recruits neural activation of multiple body parts simultaneously. Moreover, it is unclear whether various somatosensory percepts of different body parts are successfully elicited by spatiotemporally multiplexed stimulations on the brain.

Controlling the quality of artificial somatosensation is another critical issue in sensory research (Armenta Salas et al., 2018). Studies in non-human primates (NHPs) revealed that stimulus parameters such as frequency and amplitude can affect the behavioral responses of NHPs (Callier et al., 2020; Kim et al., 2015; Romo et al., 2000; Romo et al., 1998). However, NHPs could not describe the quality of sensation, such as pressure and vibration. ICMS and DCS in humans elicited various qualities of sensations during stimulation (Flesher et al., 2016; Hiremath et al., 2017; Hughes et al., 2021). Additionally, a recent human study suggested that the amplitude of ICMS current affects the quality (tactile or proprioception) of the elicited sensation (Armenta Salas et al., 2018). To date, however, fine and consistent control of elicited tactile qualities, including vibration and pressure on the same body part, remains a challenge both in ICMS and DCS studies in humans (Armenta Salas et al., 2018; Hiremath et al., 2017; Hughes et al., 2021; Lee et al., 2018).

In our previous study, we showed that downstream areas of somatosensory network including ventral premotor cortex (vPM) also elicit somatosensory percept of hand by cortical stimulation (Ryun et al., 2023). Interestingly, previous findings indicated that negative motor responses (complete inhibition of movement or a phenomenon in which the subject is unable to move by DCS even though the subject wants to move) are induced by stimulating the precentral cortex and surrounding areas, including ventral premotor cortex (negative motor phenomenon) (Filevich et al., 2012; Luders et al., 1995; Penfield and Jasper, 1954; Rech et al., 2019). Additionally, in NHPs studies, ventral premotor area (area F5) has been considered a critical region of hand movement, including grasping (Coude et al., 2019; Jeannerod et al., 1995). However, it is unclear how these findings (artificial percept for feedback, negative motor response, and critical roles in hand movement) are functionally related.

In this study, we address these three aforementioned issues using macro-level multi-site DCS and stimulation on the downstream cortical areas of the primary somatosensory cortex (S1). We showed that multi-site DCS can elicit simultaneous and independent somatosensation. We also found that multi-site DCS in small cortical regions can induce different qualities of somatosensation on the same body part. Finally, we found that DCS on vPM elicits somatosensation and negative motor response of hand simultaneously, suggesting that function-specific sensorimotor interactions exist such as hand grasping in the vPM.

## Methods

### Patient

Twenty-two patients with drug-resistant epilepsy participated in this study. Among twenty-two patients, ten patients were excluded because they had no somatosensory-related responses to stimuli or had a seizure right after stimulation. Patients underwent implantation of subdural electrode grids/strips (conventional ECoG: electrode spacing, 10 mm; diameter, 4 mm) for epilepsy monitoring. High-density electrode grids (PMT Corp., MN, USA; inter-electrode distance, 5 mm; diameter, 2 mm) were inserted on the somatosensory area in eight patients (Patients 1, 2, 6, 7, 8, 10, 11 and 12). The location of electrodes was determined based on clinical purpose. Pre-operative magnetic resonance (MR) and postoperative computed tomography (CT) images were obtained from each patient. Co-registration of the MR and CT images was performed using CURRY software (version 8.0 or 9.0; Compumedics Neuroscan) to obtain electrode locations. Functional cortical mapping with DCS for clinical purposes was performed prior to the experiment. All experimental procedures were approved by the Institutional Review Board of Seoul National University Hospital (1610-133-803 & 1612-110-816). All patients provided written informed consent before participation.

### Direct Cortical Stimulation

Stimuli were delivered through S12 and S12X cortical stimulators (Natus, Warwick, RI, USA) with charge-balanced, bipolar and constant current stimulation. The method of dynamic frequency cortical stimulation has previously been described in detail (Ryun et al., 2021). Briefly, we designed temporally varying stimulus trains using S12 cortical stimulator based on the S1 activation pattern of high-gamma activity for mechanical stimuli. That is, stimulus frequencies were continuously changed within the frequency range from 5 to 50 Hz (e.g., 50-10-20-10 Hz in 3 s). The pulse width was 0.3 ms for all stimuli. Stimulator control was performed by custom-made software written in MATLAB. To deliver stimulation triggers to the cortical stimulators, we used a 4-channel analog output, NI-9171 (National Instruments, Austin, TX, USA). To perform multi-site cortical stimulation, we used both clinically approved stimulators, S12 and S12X. During multi-site stimulation, the maximum amplitudes of the two stimulators (S12 and S12X) were 10 mA and 6 mA, respectively. The stimulation delay between two stimulators was 50 ms for two-channel multi-site stimulation. We also designed three-channel multi-site cortical stimulation using two S12 cortical stimulators and one S12X cortical stimulator. Stimulation targets were based on cortical mapping for clinical purposes and electrocorticography (ECoG) results during mechanical stimuli on the index finger. The time interval between each stimulus was at least 5 seconds for safety reasons. Additionally, ECoG data were recorded during cortical stimulation for further analysis. For Patient 1, we could not perform multi-site cortical stimulation because the patient had a seizure during single-site cortical stimulation. For Patients 2 and 3, we performed multi-site stimulation, but the second targets (the first targets were the S1) were parietal or frontal area except the S1. For Patient 4, we delivered multi-site cortical stimulation to the S1 and parieto-occipital areas. For Patient 5, we only stimulated the ventral premotor area because no electrodes were on the S1.

### Tasks

Patients were asked to report freely about their feelings during cortical stimulation. To estimate the response onset and offset time roughly, we instructed some patients to raise their hand (ipsilateral to the electrode location) when they perceived somatosensation and lower it when the sensation disappeared. We determined the response time by careful visual inspection.

For Patient 5, Ramp-and-hold pressure stimulation was manually delivered to the contralateral index finger using a von Frey tactile filament (300 g). Stimulus duration was approximately 1 s. This patient performed various upper limb movements, including grasping and reaching during DCS. This patient also performed the reach-and-grasp movement imagery task immediately after the actual reach-and-grasp movement at the same pace. This task has previously been described in detail (Jang et al., 2022).

For Patient 6, 7, and 8, DCS-guided reach-and-grasp task was performed. Patients were blindfolded and then performed a task to reach and grasp a target (arbitrary left and right position) in front of the patients. We instructed patients to: If DCS is delivered to target 1, patients move their hands one step (approximately 10 cm) to the left. Likewise, if DCS is delivered to target 2, patients move their hands one step to the right. If multi-site DCS is delivered, patients grasp the target by moving their hands forward. We calculated the accuracies of each movement (left, right and forward) and the final results (success or failure) separately.

For Patients 6, 10, 11 and 12, patients performed two alternative forced-choice tasks to determine whether changes in the qualities of somatosensation were significant. Single or multi-site cortical stimuli were delivered randomly and sequentially, and the patient selected the more pressure-like condition (for Patient 6), or the more dynamic vibration condition (for Patient 10), or the more the feeling that soft ball going down the throat condition (for Patient 11), or the more wave-like condition (for Patient 12) between the first and second (For Patient 6, single stimulation: stimulus amplitude of 2.5 mA, duration of 3 s, and frequency of 50 Hz; target 2 of multi-site stimulation: stimulus amplitude of 4 mA, duration of 2 s, and frequency of 50 Hz; for Patient 10, single stimulation: stimulus amplitude of 4 mA, duration of 2 s, and frequency of 50 Hz; target 2 of multi-site stimulation: stimulus amplitude of 4 mA, duration of 2 s, and frequency of 50 Hz; for Patient 11, single stimulation: stimulus amplitude of 5 mA, duration of 5 s, and frequency of 50 Hz; target 2 of multi-site stimulation: stimulus amplitude of 2 mA, duration of 5 s, and frequency of 20 Hz; for Patient 12, single stimulation: stimulus amplitude of 3 mA, duration of 5 s, and frequency of 10 to 50 to 10 Hz (dynamic frequency stimulation); target 2 of multi-site stimulation: stimulus amplitude of 4 mA, duration of 5 s, and frequency of 50 Hz). For Patient 10, we also delivered triple-channel cortical stimuli using two S12 and one S12X cortical stimulators.

### Data Analysis

ECoG data were recorded with the Neuroscan or Nuevo (Neuroscan, Charlotte, NC, USA) systems at 2000 Hz. ECoG channels showing epileptiform activities and abnormal fluctuations due to technical problems were excluded from further analysis. The data were re-referenced using common average reference (CAR) and notch filtered at 60 and 120 Hz. To calculate time-frequency representation, the complex continuous Morlet wavelet transform (seven cycles) was applied to the data. Transformed single-trial data were normalized by the mean and standard deviation of the resting period. We utilized linear discriminant analysis (LDA), quadratic discriminant analysis (QDA) and naïve bayes classifier for classification between reaching and grasping imagery tasks.

We used the Wilcoxon rank sum test to test the difference in response time between the onset and offset of artificial somatosensation. To calculate significance in Figure 4C, we used a nonparametric binomial test. To compare the difference in power levels of reaching and grasping movement imageries, we used a paired *t*-test.

## Results

Twelve patients with drug-resistant epilepsy were included in this study. **Figure 1** shows electrode locations in 12 patients. The demographics of 12 patients are described in **Table 1**. Before multi-site stimulation, we measured response times to stimuli and obtained behavioral reports to dynamic frequency cortical stimulation (time-varying frequency during stimulus period, see the literature (Ryun et al., 2021)). Response time of stimulus onset was 1.023 ± 0.162 s (mean ± standard error (s.e.); median = 0.648 s), and that of stimulus offset was 1.313 ± 0.3 s (median = 0.767 s) (**Fig 2**). Most patients showed fast reaction times, but some patients responded very slowly. It is presumed to be due to electrode placement or the patient’s attention level, but additional research is needed. There was no significant time difference between onset and offset (Wilcoxon rank sum test, *p* = 0.4054).

**Figure 1.**
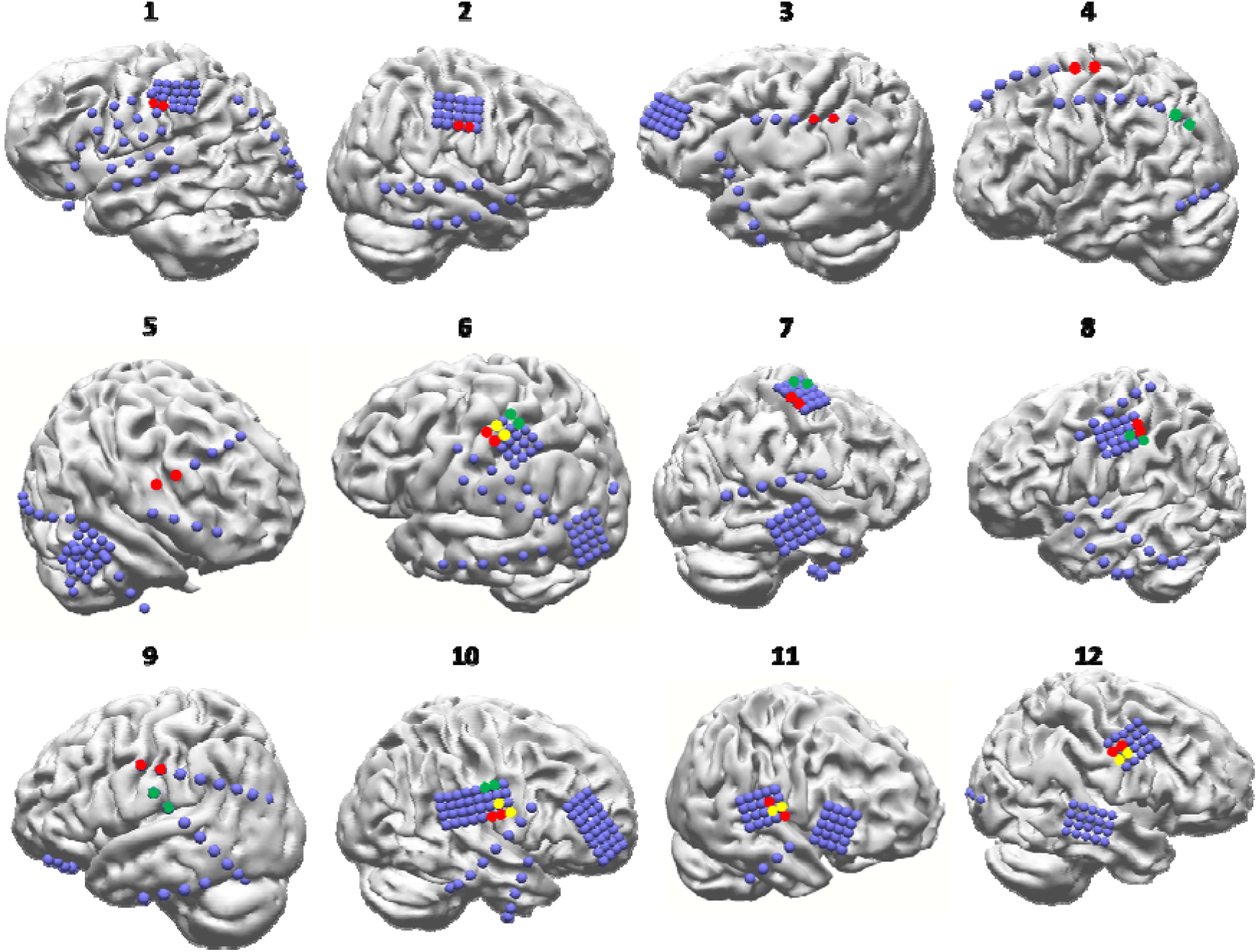
Electrode locations of all patients. Blue spheres indicate the inserted ECoG electrodes. Red circle pairs indicate the main target areas. Green circles indicate the second target areas. Yellow circles denote the second electrode pair used for changing sensation qualities. Electrode pairs were located on the S1 except for the second target of patient 4 (parieto-occipital area) and the first target of patient 5 (vPM).

**Figure 2.**
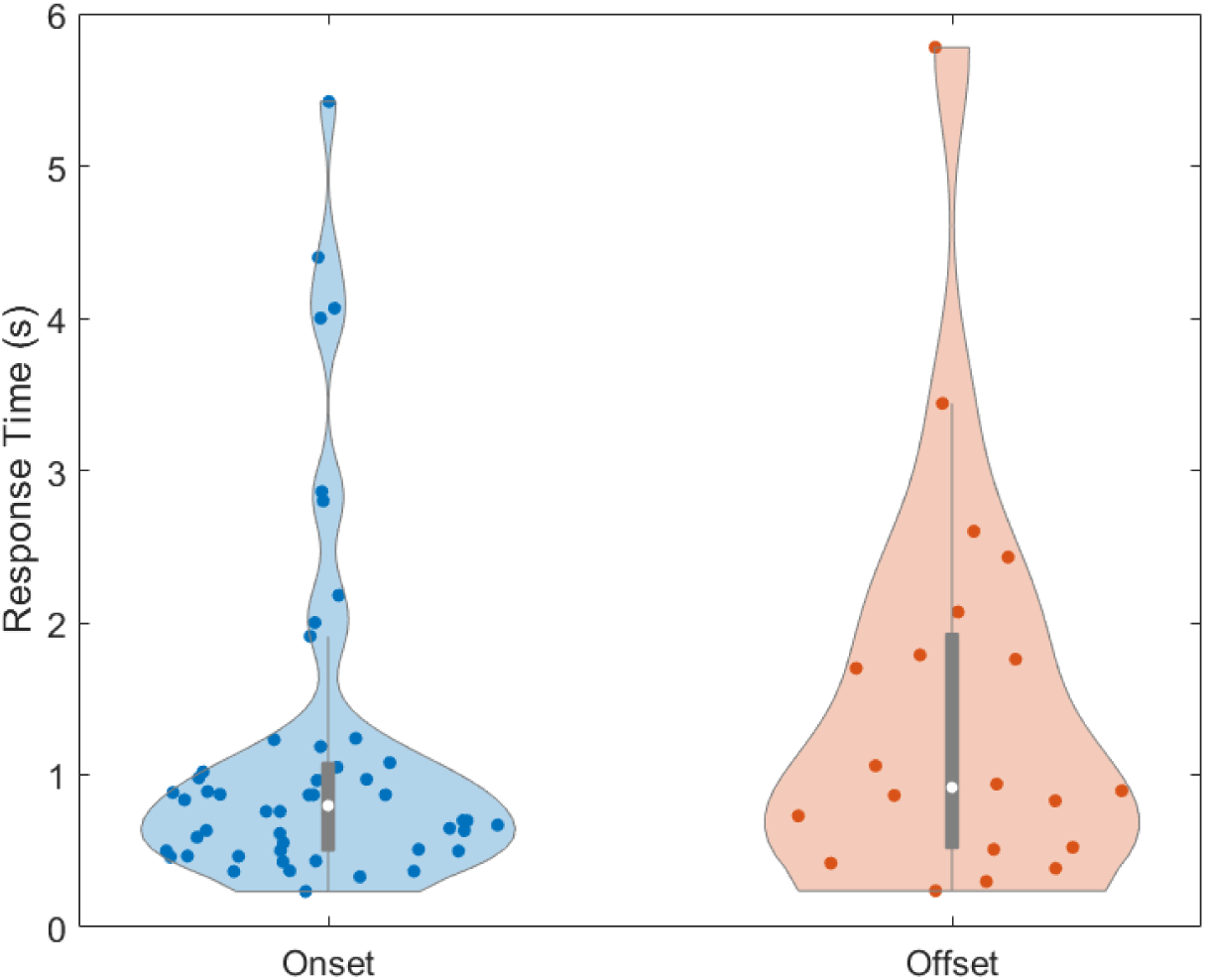
Response times of the artificial somatosensation at onset (left) and offset (right). White circles indicate the median values of each condition.

**Table 1.** Demographics of the patients.

| Patient No. | Age | Sex | Number of Electrodes | Task | Electrodes location | Diagnosis |
| --- | --- | --- | --- | --- | --- | --- |
| 1 | 40 | M | 62 | single-site DCS | Left | Hemispheric epilepsy |
| 2 | 41 | M | 40 | single-site DCS, multi-site DCS | Right | TLE |
| 3 | 29 | M | 36 | single-site DCS | Left | FLE |
| 4 | 32 | F | 32 | single-site DCS, multi-site DCS | Left | TLE |
| 5 | 21 | F | 48 | single-site DCS | Right | FLE |
| 6 | 24 | M | 60 | single-site DCS, multi-site DCS | Left | FLE |
| 7 | 29 | F | 54 | single-site DCS, multi-site DCS | Right | TLE |
| 8 | 58 | M | 40 | single-site DCS, multi-site DCS | Left | TLE |
| 9 | 31 | F | 34 | single-site DCS, multi-site DCS | Right | TLE |
| 10 | 39 | M | 84 | single-site DCS, multi-site DCS | Right | TLE |
| 11 | 30 | F | 68 | single-site DCS, multi-site DCS | Right | TLE |
| 12 | 32 | M | 68 | single-site DCS, multi-site DCS | Right | TLE |

Abbreviation: M = male, F = female, TLE = temporal lobe epilepsy, FLE = frontal lobe epilepsy.

In terms of dynamic frequency cortical stimulation (stimulus frequencies change continuously during a stimulus train), there were differences in behavioral reports between the types of inserted electrode grids. Patients with conventional ECoG electrodes implanted generally perceived the change in stimulus frequency as an immediate change in stimulus intensity (e.g., slow-fast-slow frequency dynamics = weak-strong-weak stimulus intensity) (Patient 3’s report (conventional ECoG electrode; fast (50 Hz) to slow (10 Hz) frequency vs. slow (10 Hz)-fast (50 Hz)-slow (10 Hz) frequency): *The initial sensation was pretty intense initially, but it dwindled as time went on. On the other hand, the second one seems to be getting more intense, but the numbness has not changed much.*) (Patient 4’s report (conventional ECoG electrode; fast (50 Hz) to slow (10 Hz) frequency vs. slow (10 Hz)-fast (50 Hz)-slow (10 Hz) frequency): *The sensation in my index finger started to weaken gradually with time for the first one. As for the second sensation, it gradually became stronger and spread from my second finger all the way to my thumb.*), similar to the previous study (Johnson et al., 2013). In contrast, patients with high-density ECoG electrodes implanted often perceived the change in stimulus frequency as a change in the stimulus pattern (e.g., Patient 6’s report (fast (50 Hz) to slow (10 Hz) frequency vs. slow (10 Hz)-fast (50 Hz)-slow (10 Hz) frequency): *There was a distinction between them. The first sensation was like a series of multiple thumps in my thumb, while the second one felt more like a single thump. They are definitely not the same.*) (Patient 8’s report (50 Hz vs. slow (10 Hz) to fast (50 Hz) frequency): *The sensations were distinct. The first one felt like a wave, almost like a sine wave, with a smooth and regular pattern. In contrast, the second one had a different quality to it, more like an irregular and rough wave.*). However, further research is required to generalize the difference between conventional and high-density grids. Additionally, it seemed that patients with high-density electrode grid implantation generally felt more natural sensation than patients with conventional ECoG grid implantation (**Table 2**).

**Table 2.** Receptive fields and description of sensation qualities of target areas.

| Patient No. | Electrode Type (High-density/Conventional) | Receptive Fields | Description |
| --- | --- | --- | --- |
| 1 | High-density | Tip of the right thumb | “pressure” or “twitching” |
| 2 | High-density | Left lip | “tugging” |
| 3 | Conventional | Tip of the right thumb | “tingling” |
| 4 | Conventional | Right thumb and right vision | “tingling”<br>“flickering” (vision) |
| 5 | Conventional | Left hand (palm),<br>Negative motor response | “tingling” |
| 6 | High-density | Right index finger,<br>Right little finger | “vibration”, “itching”, “pressure”<br>“pressure” |
| 7 | High-density | Left index/middle finger,<br>ulnar side of left palm | “like fine wave”<br>“tremor” |
| 8 | High-density | Right index finger,<br>right ring finger | “vibration” (not tingling)<br>“vibration” |
| 9 | Conventional | Right teeth,<br>Right lips | “like the fang moving forward”<br>“buzzing” |
| 10 | High-density | Left tongue and left lips | “itching”, “vibration” |
| 11 | High-density | Left tongue, palate and<br>throat | “pressure” |
| 12 | High-density | Left lip and mouth | “tingling”, “wave-like sensation” |

### Multi-site cortical stimulation

Multi-site cortical stimulation was performed using two or three clinically approved stimulators (S12 and S12X, Natus, Warwick, RI, USA). We performed a conventional cortical stimulation mapping procedure prior to the multi-site stimulation to determine the responsive electrodes. During the cortical stimulation mapping, 1 to 10 mA of bipolar stimuli with 1 mA step were delivered to the electrode pair located on the somatosensory-related cortical areas (stimulus duration of 5 s, frequency of 50 Hz, and pulse width of 0.3 ms). We also referred to the result of mechanical vibrotactile stimulation on the same patient and selected cortical areas which showed robust high-gamma activity as target areas for cortical stimulation.

Initially, in one patient, we found that artificial somatosensation and visual flickering are simultaneously elicited during multi-site cortical stimulation (Patient 4 in **Fig 1**). The patient reported a tingling sense in the right thumb with flickering in the right visual field (Stimulation 1 (MNI coordinates: -41, -24, 62): stimulus frequency of 50 Hz, amplitude of 5 mA (tingling sense); Stimulation 2 (MNI coordinates: -52, -65, 18): 50 Hz, 4 mA (flickering)). That is, multi-site DCS could elicit different sensory modalities (somatosensory and vision) simultaneously. We applied this phenomenon to the same sensory modality based on this result. We performed simultaneous cortical stimulation in various S1 regions for five patients to elicit simultaneous multiple artificial somatosensations. All five patients reported simultaneous and independent somatosensory senses in different body parts. In the case of Patient 6, the patient reported a vibration sense in the radial side of the index finger when stimulating the ventral area of the S1 (MNI coordinates: -50, -26, 59; high-density electrode; 50 Hz, 2.5 mA) and a pressure sense in the first knuckle of the little finger when stimulating the relatively dorsal area of S1 (MNI coordinates: -38, -36, 67; high-density electrode; 50 Hz, 1.5 mA). When both cortical areas were stimulated, the patient felt the vibration sense in the index finger (Patient 6’s report: *I am experiencing a sensation… of either vibration or itching in my index finger…*) and the pressure sense in the little finger (Patient 6’s report: *It is like someone is pressing on my little finger with their hand.*) simultaneously, without any distortion of the sense due to the simultaneous stimulation itself.

Given this result, we designed an artificial somatosensory feedback-based reach-and-grasp task. Three patients participated in this task. For Patient 6, we asked the patient to move his arm to the left if he felt sensation in the radial side of the index finger (stimulus amplitude of 2.5 mA, duration of 3 s, and frequency of 50 Hz), and to the right if he felt sensation in the little finger (stimulus amplitude of 1.5 mA, duration of 2 s, and frequency of 50 Hz). When he felt both, he moved his arm forward, and if there was an object there, he grabbed it. Blindfolded patients could efficiently perform the task with a high success rate (**Fig 3** and **Movie 1**). Success rates of each movement performance (left, right and reach-and-grasp movements) were 100% (52/52; success trials/total trials), 92.5% (74/80) and 91.4% (74/81) in Patients 6, 7, and 8, respectively. The success rates of the object grasping task were 100% (17/17), 75% (18/24), and 94.7% (18/19) in Patients 6, 7 and 8, respectively (**Fig 3**). Patient 7 felt sensations like “fine wave” in the index and middle finger (MNI coordinates: 41, -29, 60; high-density electrode; stimulus amplitude of 5 mA, duration of 3 s, and frequency of 50 Hz) and felt tremor or vibration sense in the ulnar side of the palm (Patient 7’s report: *I experienced a trembling sensation in the palm on the side closer to the pinky finger of my left hand. It is similar to the feeling of a golf ball bouncing back and forth in my palm.)* (MNI coordinates: 28, -30, 70; high-density electrode; stimulus amplitude: 4 mA, duration: 2 s, frequency: 50 Hz). Patient 8 felt vibration senses both in the index (MNI coordinates: -61, -23, 46; high-density electrode; stimulus amplitude: 5 mA, duration: 3 s, frequency: 50 Hz) and ring fingers (MNI coordinates: -60, -28, 49; high-density electrode; stimulus amplitude: 7 or 7.5 mA, duration: 2 s, frequency: 50 Hz) (**Table 2**). Additionally, one patient (Patient 9) performed three-alternative forced-choice task. The patient was instructed to determine where the sensation is elicited among the lip (Patient 9’s report: *I have this sensation where it feels like my lower lip is buzzing or vibrating.*) (MNI coordinates: -60, -15, 38), tooth (Patient 9’s report: *It is almost like my front tooth, the fang, is shifting forward.*) (MNI coordinates: -67, -19, 21) and both sides during single or multi-site cortical stimulation (success rate = 62.9% (22/35), chance level = 33.3%). For Patient 10, the patient felt sensations in three different sites: the left side of the face, the tip of the tongue, and the left side of the tongue. We designed four alternative, forced-choice tasks using three cortical stimulators simultaneously. We delivered four different stimuli to the responsive electrode pairs (e.g., target 1, target 2, target 3 and target 1-2). The success rate of behavioral performance was 81.13% (43/53, chance level = 25%).

**Figure 3.**
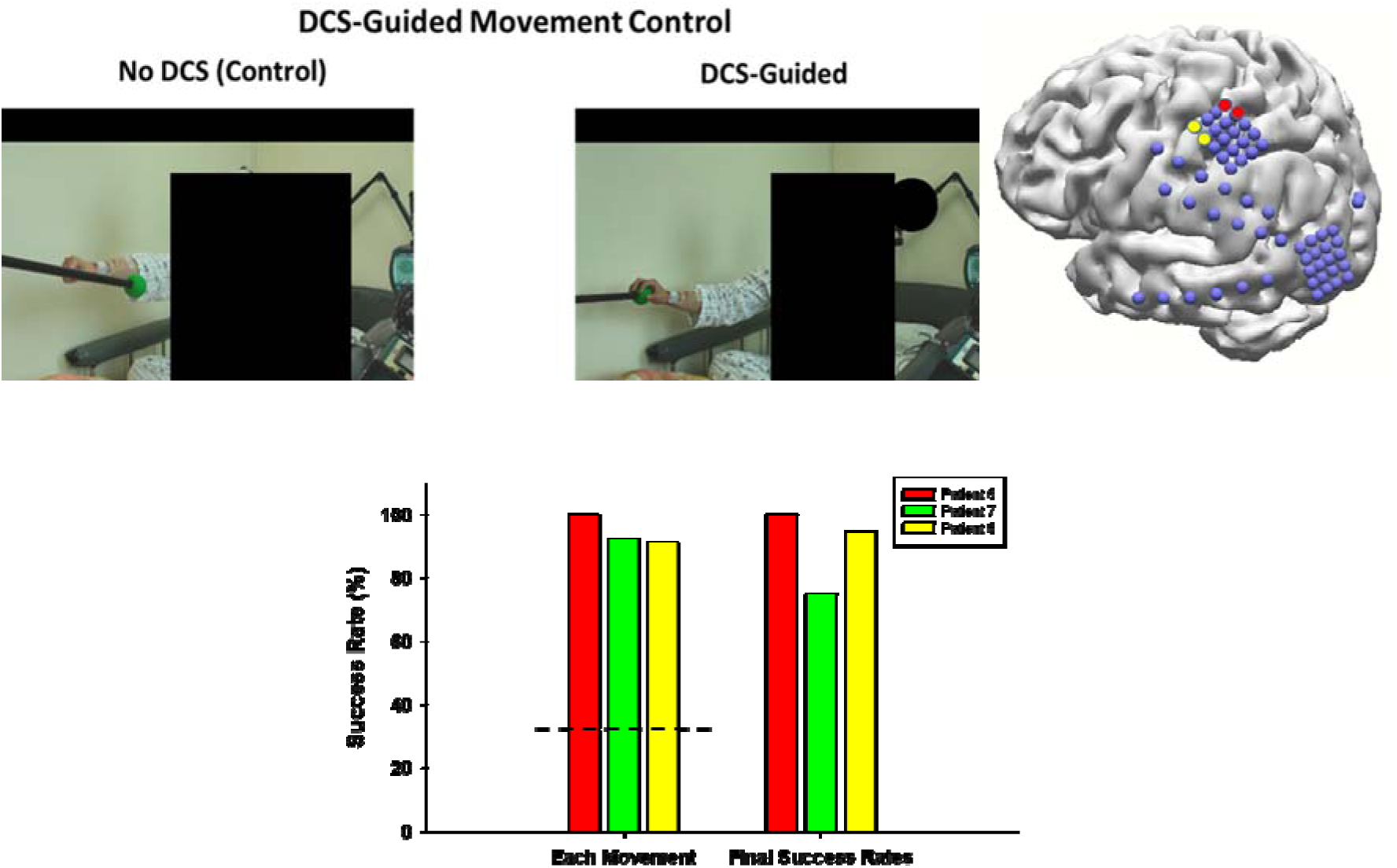
(Top left) Movement control (reach-and-grasp) by multi-site DCS. Patient 6 was instructed to move his arm left when he feels the sensation in the index finger, move right when he feels the sensation in the little finger, and move his arm forward and grasp when he feels both sensations (snapshots from the video results). For safety reasons, the time interval between each stimulus was at least 5 seconds. (Top right) Location of stimulating electrode pairs of Patient 6. (Bottom) Success rates of each movement (left, right, and reach-and-grasp motions) performance and final (grasping the target) success rates. Dashed horizontal line indicates chance level (33 %).

**Figure 4.**
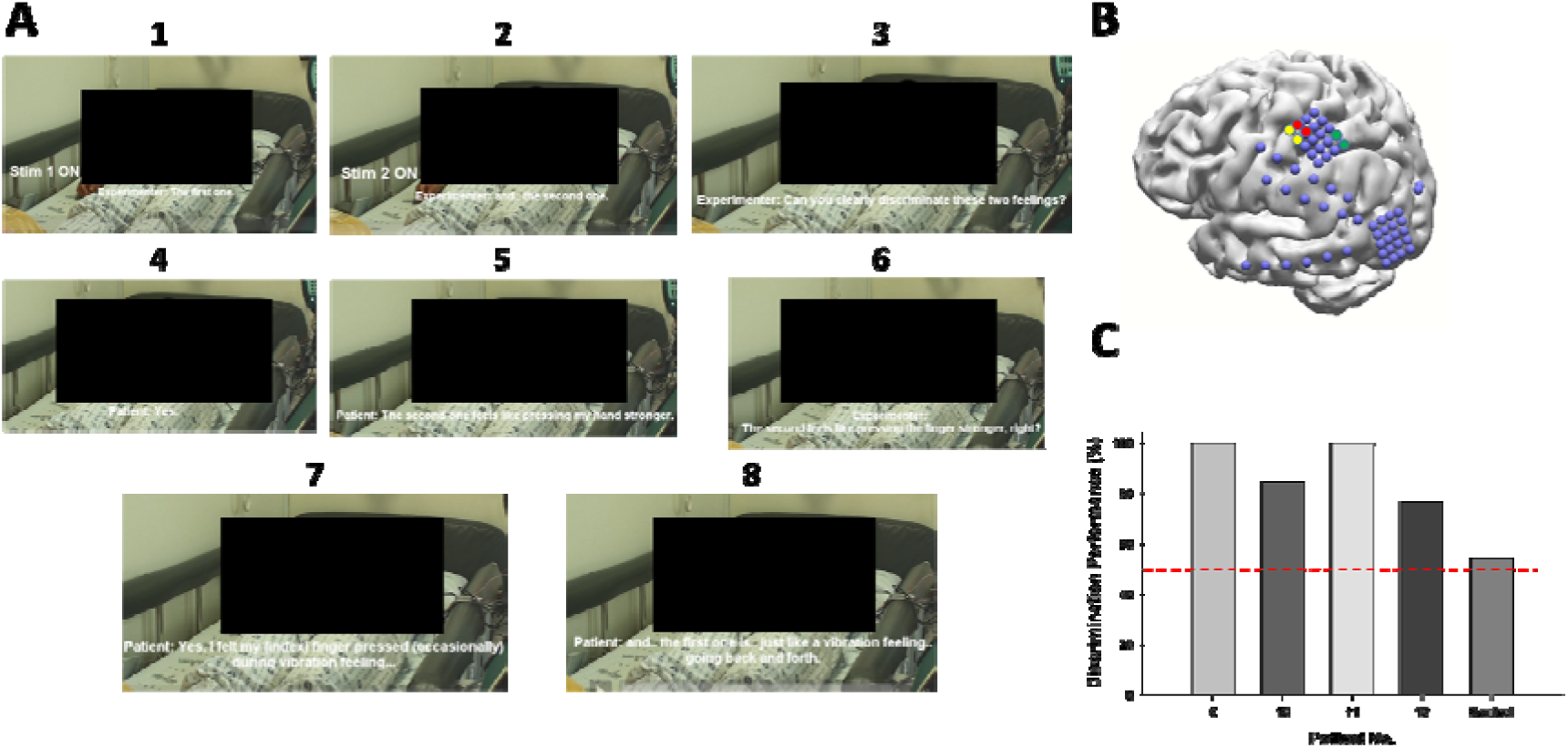
(A) Multi-site DCS on the areas close to each other. The first stimulus was the single-site DCS, and the second one was the multi-site DCS, including the area of the first stimulus. Numbers above the snapshots indicate the sequence of events. (B) Electrode locations of the representative patient (Patient 6). Yellow and red electrode pairs were used for multi-site cortical stimulation. (C) Discrimination performance of the two-alternative forced-choice task in four patients. Four left bars indicate the condition that two electrode pairs are close to each other, and the control indicates the condition that one electrode pair is located on the S1 (yellow) and the other electrode pair (green) is located on the other parietal area where the area expected to have little relevance to sensory perception in Patient 6 (n =11). Discrimination performances of four patients were significantly high (nonparametric binomial test, *p* = in Patient 6 (n = 21), *p* = in Patient 10 (n = 35), *p* = in Patent 11 (n = 21), and *p* = 0.0026 in Patient 12 (n = 30)).

Interestingly, we also found that multi-site cortical stimulation on S1 area close to each other induces different quality of somatosensation in the same body part from 4 patients. In Patient 6, one single-site DCS on S1 (MNI coordinates: -50, -26, 59; high-density electrode; 50 Hz) elicited vibration sense on the radial side of the index finger, and the other one (MNI coordinates: -46, -29, 61; high-density electrode; 20 Hz) elicited the same quality of sensation (vibration sense) on the ulnar side of the index finger. However, DCS on both areas elicited vibration and pressure senses on the radial side of the index finger, but no sensation was elicited on the ulnar side of the index finger (**Fig 4** and **Movie 2**). To confirm the consistency of this phenomenon, the patient performed two alternative, forced-choice task between single (elicited sensation: vibration sense on the radial side of the index finger) and multi-site (elicited sensation: vibration and pressure senses on the radial side of the index finger) cortical stimulation. That is, two (single and multi-site DCS) stimuli were delivered randomly and sequentially, and the patient selected a condition that felt more pressure-like between the first and second one. The accuracy of this task was 100% (21/21; **Fig 4**). In Patient 10, one single-site DCS on S1 (MNI coordinates: 64, -10, 23; high-density electrode; 50 Hz) elicited vibration sense on the tip of the tongue, and the other one (MNI coordinates: 66, -11, 20; high-density electrode; 50 Hz) elicited the same sensation on the left tongue. When we delivered stimulation to both electrode pairs, the patient felt it as a vibration sense with spatiotemporal dynamics (e.g., vibration sense from the left side of the tongue to the top of the tongue). This patient also performed two alternative, forced-choice tasks between single and multi-site stimulation conditions. The accuracy of the task was 84.38% (27/35). In Patient 11, one single-site DCS (MNI coordinates: -65, -15, 18; high-density electrode; 50 Hz) elicited pressure sense on the tongue and palate, but the other one (MNI coordinates: -65, -14, 15; high-density electrode; 20 Hz) elicited no sensation. When we delivered stimulation to both electrode pairs, the patient reported feeling like a soft ball going down her throat (accuracy: 100% (21/21)). In Patient 12, one single-site DCS (MNI coordinates: 62, -12, 37; high-density electrode; slow (10 Hz)-fast (50 Hz)-slow (10 Hz)) elicited tingling sense on the medial side of the left lip, and the other one (MNI coordinates: 62, -10, 35; high-density electrode; 50 Hz) elicited the same sense on the lateral side of the left lip. When we delivered stimulation to both electrode pairs, the patient reported a wave-like tingling sensation on the medial side of the lip only (accuracy: 76.67% (23/30)) (Fig. 4C).

### Cortical stimulation on vPM

Next, we tested the hypothesis that the three independent findings mentioned earlier ((i) artificial somatosensation, (ii) negative motor response during vPM stimulations, and (iii) findings from NHP studies indicating the crucial role of vPM in hand grasping) are functionally related. We first tried to elicit artificial somatosensation in the vPM area. In Patient 5, the patient consistently reported a tingling sense in the contralateral palm, including all fingers, consistent with our previous finding (Ryun et al., 2023). We also found strong high-gamma activity in this area (MNI coordinates: 51, 7, 40) during mechanical pressure stimulation on the index finger, consistent with previous findings (Avanzini et al., 2016; Ryun et al., 2023). In the main experiment, we asked the patient to perform various upper limb movements, including reaching and grasping during DCS. Interestingly, we observed negative motor responses only when the patient performed grasping motions (**Fig 5** and **Movie 3**). In other words, the patient was unable to grasp the target even though she had the will to grab it while feeling a consistent tingling in her palm. No abnormalities, including muscle contraction (except palmer sensation), were observed when DCS was delivered at rest. No significant movement abnormalities were found during elbow flexion and reaching without grasping (**Movie 3**). The patient also performed a reach-and-grasp movement imagery task. Interestingly, strong high-gamma activity and alpha/beta event-related desynchronization (ERD) were found during the grasping imagery period, compared to those during the reaching imagery period (**Fig 5D**). Classification accuracy between these reaching and grasping imagery periods was 72.6 to 75 % (**Fig 5E**). These results suggest that vPM is a core region for sensorimotor interaction and planning or imagination about hand movement.

**Figure 5.**
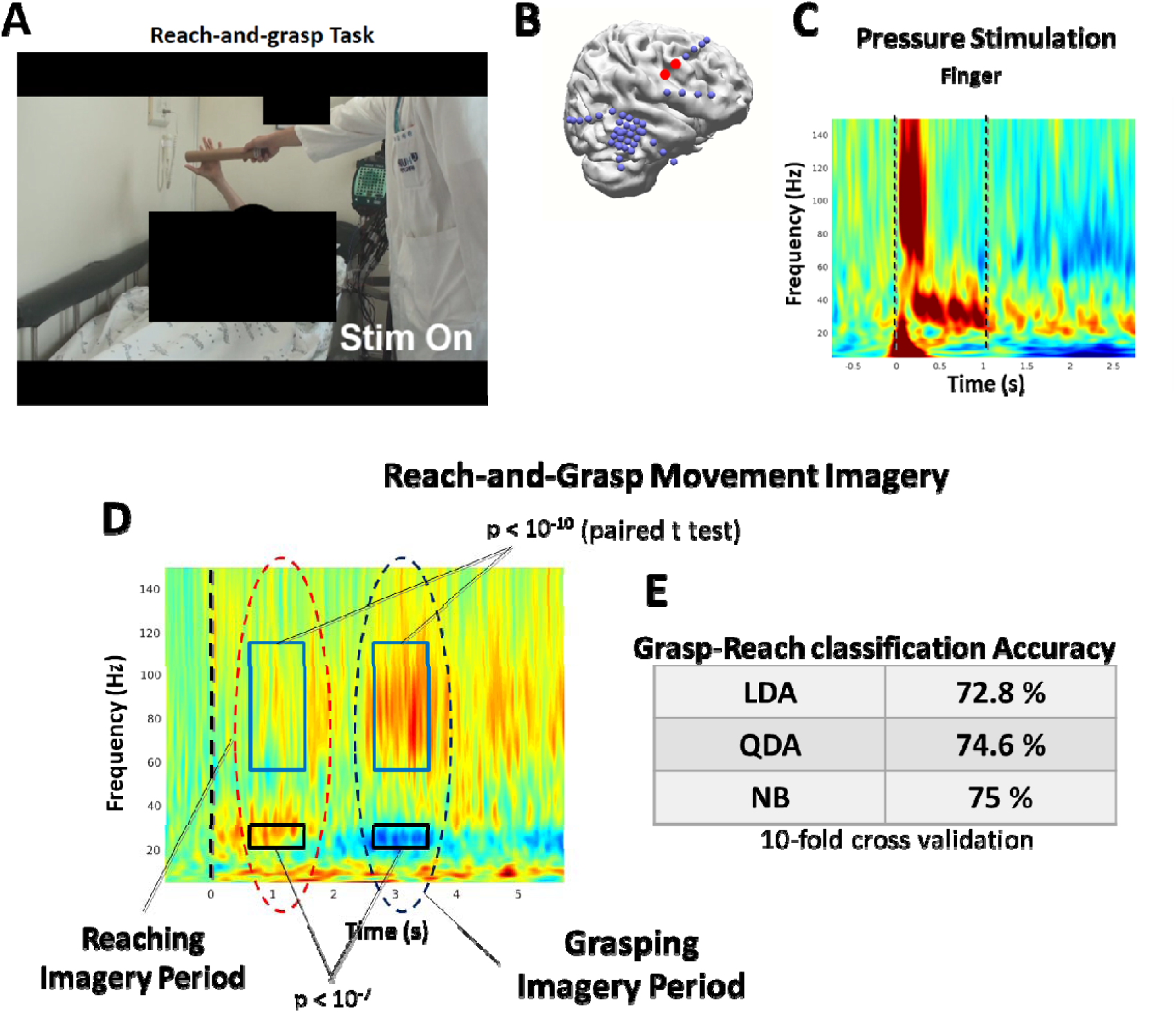
(A) Behavioral responses during DCS on vPM. DCS elicited artificial somatosensory perception of the hand (palm side) and negative motor response of the hand simultaneously. (B) Electrode location. (C) Time-frequency representation of vPM area during mechanical pressure stimulation. Dashed vertical lines indicate stimulus onset and offset. (D) Time-frequency plot during reach-and-grasp imagery and (E) classification accuracy between reaching and grasping. The dashed vertical line at t = 0 indicates the onset of the imagery period presenting a target. (abbreviation: LDA = linear discriminant analysis, QDA = quadratic discriminant analysis, NB = naïve Bayes classifier)

## Discussion

In this study, we performed multi-site DCS and DCS on the sensorimotor downstream area to address three questions: (i) Can the multi-site DCS replicate the spatiotemporal dynamics of somatosensory feedback and (ii) control the quality of elicited somatosensation, and (iii) how and where does the neural feedback activity interact with neural activity for motor processing? We found that multi-site DCS on the S1 can elicit simultaneous and independent tactile sensations and confirmed that its robustness is sufficient to apply the somatosensory feedback system by performing DCS-guided movement control task. We also found that multi-site DCS close to each other in the S1 can change the quality of the tactile sensation (e.g., vibration and pressure) on the same body part. Additionally, we found that DCS on the vPM can elicit artificial somatosensation and negative motor response simultaneously and inhibits grasping motion selectively in reach-and-grasp task while eliciting consistent somatosensation in the palm, which is an essential body part for somatosensory feedback during hand grasping. This result might reflect a function-specific (e.g., grasping) sensorimotor interaction in the vPM.

### Replicate the spatiotemporal dynamics of somatosensory feedback

Our results showed that multi-site DCS can elicit sensations independent of each other and temporally stable. These characteristics enable the generation of somatosensory feedback that is dynamic in both space and time with a high degree of freedom. In terms of BMI applications, although the best scenario of the somatosensory feedback is to replicate the exact feedback of evoked sensation during limb movement or object manipulation, it may be challenging to practically apply to the BMI system because of the limited spatial coverage of the electrodes both at the macro and micro levels. An alternative approach is to match the event with a specific elicited artificial sensation. This type of feedback system has been performed in monkey and rodent studies (O’Doherty et al., 2011; Thomson et al., 2013; Venkatraman and Carmena, 2011). In the current study, we showed that such an alternative approach works well throughout the experiment with blindfolded patients. Although there is an unavoidable mismatch between “real” and alternative somatosensory representations, this learning-based feedback approach may be helpful if it is impossible to elicit exact somatosensory feedback. In the dynamic frequency cortical stimulation experiment, behavioral reports after stimulation were different depending on the type of electrodes (conventional vs. high-density electrode). Generally, increasing stimulus frequencies increased the intensity of elicited somatosensation. However, in some cases, dynamic frequency stimulation by high-density electrodes did not induce an increase in somatosensation intensity but changed the pattern of elicited somatosensation. This result indicates that more focal stimulation with frequency dynamics can control the detail of elicited somatosensation, although it is not exactly known how focal stimulation is needed to control the sensation detail. It could be explained by the difference in the diameter of the electrode, which may affect the current spread of stimulation, and the stimulation depth, which determines the cortical layers stimulated. Further research, including modeling approach, is needed to quantify the effects of these stimulation conditions.

### Change of sensation quality by multi-site DCS

Another interesting result is that multi-site DCS on the S1 area close to each other elicited different qualities of somatosensation in the same body part. Perhaps there are two possible scenarios to interpret this result. The first one is electric field modulation due to the multi-site cortical stimulation. Although the charge density is highest at the electrode site, subtle changes in the electric field at the surrounding area induced by the current of each electrode pair may induce the change in sensation quality. Moreover, since DCS was delivered with different stimulation parameters (e.g., 20 Hz and 50 Hz) to each cortical region, the spatial distribution generated by electrical stimulation is different depending on the type of stimulation (unipolar vs. bipolar) (Nathan et al., 1993). Additionally, stimulus parameters, including inter-electrode distance, stimulus amplitude and frequency, can all affect the current density distributions (Fiocchi et al., 2018). The second possible scenario is that the quality of somatosensation may be built by the spatial combination of neuronal activity in the S1. Indeed, although body representation follows homunculus, the representing areas are not clustered strictly but locally mixed across the areas 1 and 2 in the S1 (Iwamura et al., 1980; Janko et al., 2022; Kurth et al., 2000). Expressly, several studies indicated a mosaic organization in S1 (Fardo et al., 2018; Favorov and Diamond, 1990; Favorov et al., 1987). Although we do not know the actual mechanism at this point, it is quite clear that including the spatiotemporal dynamics of neural activity is essential for eliciting various qualities of somatosensation. In this respect, multi-site brain stimulation with spatiotemporal dynamics can be a promising technique to replicate the complex somatosensory feedback artificially.

### Functional role of vPM area

Our vPM results indicate that this area is important for both somatosensation and action. In a monkey study, it is known that vPM is critically involved in the somatosensory-to-action transformation (Romo et al., 2004). Additionally, neurons in the vPM were activated during passive somatosensory stimulation in monkey and human studies (Avanzini et al., 2016; Rizzolatti et al., 1981). That is, the vPM is involved in the bottom-up sensorimotor processing. On the other hand, a human study suggested that this area modulates the S1 during voluntary movement without proprioceptive feedback (Christensen et al., 2007). We observed a robust high-gamma activity and alpha/beta ERD during the grasping imagery task. In light of these findings, it is thought that vPM area is a crucial region for both bottom-up and top-down sensorimotor processing.

In the present study, we found that the body part that shows both negative motor response and artificial somatosensation is restricted to the hand only. Many previous studies have indicated that vPM is involved in the planning/execution of hand grasping/object manipulation by visuomotor transformation (Fogassi et al., 2001; Hoshi and Tanji, 2007; Murata et al., 1997). Our result shows that vPM is involved not only in the visuomotor transformation and movement planning, but also in the sensorimotor integration for hand movements. In light of these findings, the vPM area may be a cortical hub for hand movement throughout the multisensory-motor integration, including vision and somatosensation.

The present study suggests that somatosensory downstream areas such as vPM can induce artificial somatosensation with specific motor inhibition in the same body part. It is unclear whether the stimulation on the vPM evokes artificial somatosensation by activating the S1 through the fiber tract or it is induced by the activation of vPM only. Suppose the latter is true (artificial sensation was independent of the S1). In that case, it demonstrates that somatosensation can be induced without the S1 region, and thus, it may be helpful for restoring somatosensory function for people with lesions in the S1 area.

### Perspective and Limitation

A recent ICMS study suggested that biomimetic multi-channel ICMS can induce high resolution of force feedback (Greenspon et al., 2023). However, multi-channel ICMS stimulating a very narrow (hundreds of μm spacing) area did not change the quality of elicited somatosensation. Given this finding, our results indicate that large-scale multi-site cortical stimulation can be a promising approach to finely control the elicited somatosensation. Indeed, previous studies have suggested that there are several cortical regions where artificial somatosensations are elicited by the DCS (Balestrini et al., 2015; Caruana et al., 2018; Ryun et al., 2023). In Patient 8, we could not change the quality of elicited somatosensation by multi-site DCS. It is not easy to interpret because we do not exactly know the underlying neural mechanisms of DCS and neural characteristics of stimulated areas. It is probably due to the location of the two electrode pairs, but further studies, including investigation of the neural response mechanism of the DCS, are needed.

Eliciting artificial somatosensation via direct brain stimulation is critically needed in bi-directional brain-machine interfaces (BMI). It can improve the performance of robotic arm control in the BMI system and plays a vital role in inducing a sense of ownership of our body parts (Bensmaia and Miller, 2014; Collins et al., 2017; Flesher et al., 2021). Our present results may provide insight into how sensory feedback should be provided through brain stimulation in bidirectional BMI systems.

## Supporting information

Supplementary Movie 1-2 Control

## Acknowledgments

We thank Yujin Yang for performing the movement imagery experiment. This research was supported by the Alchemist Brain to X (B2X) Project funded by Ministry of Trade, Industry and Energy (20012355, NTIS: 1415181023).

## Author Contributions

S. Ryun and C. K. Chung designed and conducted the study. C. K. Chung performed the surgeries. S. Ryun designed the tactile devices and experimental paradigm. S. Ryun carried out experiments. S. Ryun and C. K. Chung performed data analysis and wrote the paper.

## Competing Financial Interests

The authors declare that there are no competing financial interests regarding the publication of this paper.

## Supplementary Information

**Supplementary Movie 1-1.** A DCS-guided reach-and-grasp task of Patient 6. Blindfolded patients performed reach-and-grasp tasks for grabbing target objects using information from multi-site DCS.

**Supplementary Movie 1-2.** Reach-and-grasp task without DCS (control condition).

**Supplementary Movie 1-3 and 4.** DCS-guided reach-and-grasp task of Patients 7 and 8.

**Supplementary Movie 2.** Patient’s report of the multi-site DCS on the S1 close to each other.

**Supplementary Movie 3.** Behavioral results of DCS on vPM.

## Notes

### Competing Interest Statement

The authors have declared no competing interest.

